# Skin microbiome composition and function in the development of atopic diseases during infancy

**DOI:** 10.64898/2025.12.22.696050

**Authors:** Zeyang Shen, Jana Eckert, Richard Saffery, Katrina J. Allen, Audrey Walsh, VITALITY team, Clay Deming, Qiong Chen, Karen Laky, Jenny Min Li, Lindsay Chatman, NISC Comparative Sequencing Program, Heidi H. Kong, Kirsten P. Perrett, Julia A. Segre, Pamela A. Frischmeyer-Guerrerio

## Abstract

**Background:** Atopic dermatitis (AD), food sensitization (FS), and food allergy (FA) frequently co-occur in infancy, but the factors driving distinct atopic phenotypes remain unclear. While *FLG* null mutations are major genetic risk factors for AD, they explain only a fraction of disease heritability, suggesting a potential role for the skin microbiome.

**Objective:** To determine how early-life skin microbiome composition and its interaction with host genetics contribute to distinct atopic phenotypes in infancy.

**Methods:** We analyzed >1,000 skin swabs from 429 infants in the VITALITY cohort using deep shotgun metagenomic sequencing at 2-3 months (pre-diagnosis) and 12 months (post-diagnosis). Differential abundance, strain-level, and microbial genome-wide association analyses were performed to identify taxonomic and functional features associated with AD, FS, FA, and their co-occurrence, as well as with *FLG* mutation status.

**Results:** Within AD, microbial signatures differed by co-occurring FA or FS. At 12 months, *Staphylococcus epidermidis* was enriched in infants with AD alone, whereas infants with AD and FA exhibited decreased *Staphylococcus hominis* and *Lactococcus* species, along with increased *Dermacoccus nishinomiyaensis* and *Malassezia slooffiae*. At 2-3 months, early skin dysbiosis characterized by enrichment of *Staphylococcus* species was associated with subsequent development of AD with FS or FA, but not AD alone. Among infants with AD, *FLG* mutation carriers exhibited additional microbial shifts, including reduced *Streptococcus* species and increased *Malassezia slooffiae*. Strain-level analyses revealed mother–infant sharing of skin microbial taxa associated with AD, and microbial genome-wide association analyses identified species-specific genes linked to AD severity.

**Conclusions:** Infant atopic phenotypes are associated with distinct, phenotype-specific features of the skin microbiome that emerge both before and after clinical disease onset. By resolving microbial differences within AD according to allergic co-occurrence, host genetics, and early-life timing, this study highlights the infant skin microbiome as a potential target for early risk stratification.

## INTRODUCTION

Allergic diseases are among the most common chronic health conditions affecting both children and adults and frequently begin in infancy^1^. Atopic dermatitis (AD) and food sensitization (FS) or food allergy (FA) often represent the earliest manifestations of the atopic march. AD affects up to 20% of children, with most cases presenting during the first year of life^2–4^. Co-occurrence of AD with FS or FA is frequent^5^, yet the factors that drive divergence in early atopic outcomes are not fully understood.

Null mutations in *FLG*, which encodes the epidermal barrier protein filaggrin, represent the strongest known genetic risk factor for AD^6,7^, and have also been associated with increased risk of FA^8^. However, known genetic factors explain only about 20% of the total heritability of AD and FA^7,8^, suggesting an important role for non-genetic contributors, including the microbiome.

The skin microbiome is altered in both children and adults with AD compared to healthy individuals, including shifts in the abundance and strain-level diversity of staphylococcal species^9–14^. Although these findings implicate skin microbes in AD pathogenesis, it remains unclear whether microbial alterations precede disease onset or arise as a consequence of skin inflammation and barrier dysfunction.

Early-life studies using 16S rRNA gene sequencing suggest that infant skin colonization patterns may influence subsequent AD risk, but results have been inconsistent, particularly with respect to *Staphylococcus aureus* colonization^15–17^. Moreover, the role of early-life skin dysbiosis in FS and FA, and its relationship to AD and allergic co-occurrence, remains poorly defined despite evidence that *S. aureus* colonization can distinguish AD with FA from AD alone and is associated with FA independent of eczema severity^18,19^.

In this study, we investigated how early-life skin microbiome composition and function contribute to the development of AD, FS, and FA during the first year of life. Using the VITALITY infant cohort^20^, we analyzed infant and maternal skin samples collected at 2–3 months and 12 months of age using deep shotgun metagenomic sequencing, enabling species-, strain-, and gene-level resolution of microbial features associated with distinct atopic phenotypes.

## METHODS

### Study design

The VITALITY trial is a randomized, double-blind, placebo-controlled study designed to assess whether oral vitamin D supplementation during infancy reduces the risk of food allergy at 12 months of age. Healthy, term infants were recruited at 6–12 weeks of age in metropolitan Melbourne, Australia. Atopic dermatitis (AD) was diagnosed by trained clinicians at 6 and/or 12 months, with disease severity assessed using the SCORAD index^21^. Food allergy (FA) was confirmed by oral food challenge, and food sensitization (FS) was defined by positive skin prick testing and/or food-specific IgE with clinical tolerance^22^. This study was conducted as a nested, phenotype-enriched microbiome analysis within the VITALITY cohort.

### Sample collection and negative controls

Skin samples were collected from the cheek and antecubital fossa at 2–3 months of age (baseline) and at 12 months. Procedures for sample collection, negative controls, and sample exclusion criteria are described in the Supplementary Methods.

### Shotgun metagenomic sequencing and processing

Shotgun metagenomic library preparation, sequencing, and read processing were performed as previously described^12,23,24^ and are detailed in the Supplementary Methods.

### Diversity and differential abundance analyses

Alpha and beta diversity analyses and differential abundance testing were conducted using established methods, as detailed in the Supplementary Methods.

### Host genetics and microbial association analyses

Host genetic variation in the filaggrin gene (*FLG*) was inferred directly from metagenomic data with stringent variant-calling criteria (see Supplementary Methods). Differential abundance analyses were conducted to identify microbial features associated with *FLG* mutation status.

### Microbial GWAS and strain tracking

Microbial genome-wide association analyses were performed to identify species-specific microbial genetic variants associated with AD severity using custom frameworks (see Supplementary Methods). Strain sharing between infants and mothers was assessed using reference-based strain tracking methods and permutation testing to evaluate statistical significance.

### Statistical analysis

Pairwise group comparisons were conducted using Mann–Whitney U tests. For analyses involving multiple hypothesis testing, p values were adjusted using the Benjamini–Hochberg procedure.

## Data and code availability

The raw metagenomic sequencing data are available in the NCBI BioProject database under project number PRJNA971252. Analysis code and pipelines are available at: https://github.com/zeyang-shen/VITALITY_skin_atopy.

## RESULTS

### Characteristics of study population

A total of 429 infants were included in this study. Among them, 95 were diagnosed with at least one IgE-mediated food allergy (FA) by 1 year of age based on oral food challenges, including 73 infants with co-occurring atopic dermatitis (AD) and 22 with FA alone. An additional 55 infants were food tolerant but classified as food sensitized (FS) based on positive skin prick testing and/or food-specific IgE, including 31 with co-occurring AD and 24 with FS alone. We also included 171 infants with AD alone and 108 healthy controls (HCs) without AD, FA, or FS.

Skin swabs were collected from the cheek and antecubital fossa at the 2–3-month baseline visit (n = 80 infants; median age, 3.4 months; IQR, 2.9–3.7 months) and at the 12-month clinic visit (n = 425 infants; median age, 13.1 months; IQR, 12.5–13.9 months) (Fig. 1a). At 12 months, 70 infants were sampled from AD-affected antecubital fossa, including 43 with AD alone, 19 with AD and FA, and 8 with AD and FS. Most infants with AD had mild disease, with a median SCORAD score of 7.9 (IQR, 7–17.4), and SCORAD did not differ significantly among AD subgroups (Fig. 1b). Additional clinical and demographic characteristics are summarized in Table E1.

**Figure 1.**
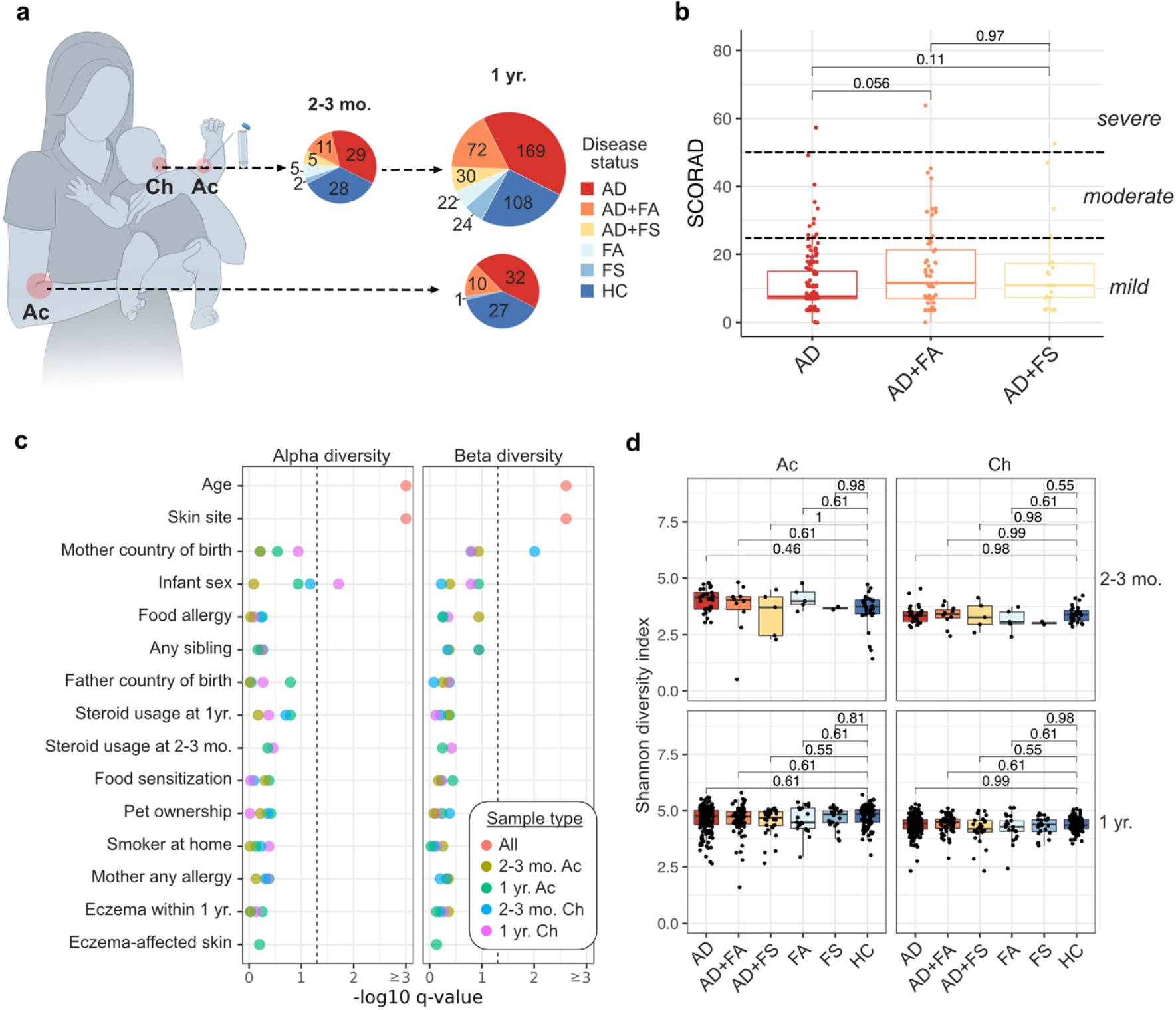
Overview of skin metagenomic data. (**a**) Summary of infants and skin swabs analyzed in this study. Modified from a related study. Numbers inside the pie chart represent the number of participants in each disease state. (**b**) SCORAD scores of infants grouped by disease status. Classification of AD severity (mild, moderate, severe) based on SCORAD is indicated. (**c**) Statistical significance of the effects of clinical and demographic variables on microbiome alpha and beta diversity. Age and skin site were tested across all samples, while the other variables were tested within groups defined by age and skin site. The dotted line indicates the significance threshold of 0.05. (**d**) Shannon diversity index. Pairwise comparisons were tested using Mann–Whitney U tests. Q values were calculated using the Benjamini–Hochberg procedure. Ac: antecubital fossa. Ch: cheek. AD: atopic dermatitis. FA: food allergy. FS: food sensitization. HC: healthy control. SCORAD: scoring atopic dermatitis.

### Overview of skin metagenomes

This study included a total of 1,078 skin samples collected from the cheek and antecubital fossa, comprising 159 samples from infants at 2–3 months of age, 849 samples from infants at 12 months of age, and 70 antecubital fossa samples from mothers collected at the 12-month visit.

All samples were analyzed using deep shotgun metagenomic sequencing.

To assign taxonomy to non-human reads, we constructed an integrated reference database comprising RefSeq genomes, two published skin microbial genome catalogs^25,26^, and 250 newly assembled infant skin metagenome-assembled genomes (MAGs) (Table E2; Fig. E1). Using this database, a median of 76% of non-human shotgun reads could be taxonomically classified (Fig. E1c).

Alpha diversity (Shannon index) and beta diversity (Bray–Curtis dissimilarity) were calculated and assessed against clinical and demographic variables (Fig. 1c). Age and skin site were the strongest determinants of microbial diversity. After stratifying by age and skin site, additional associations were observed with maternal country of birth and infant sex (Fig. E1d,e). No differences in alpha diversity were observed between disease groups (Fig. 1d) or between AD-affected and unaffected skin (Fig. E1f).

### Distinct skin microbial signatures associated with atopic diseases

We next investigated whether microbial signatures could distinguish HCs from infants with atopic disease using differential abundance analysis. At 12 months, microbial patterns varied by both skin site and allergic status. No taxa differed significantly between participants with FS or FA alone and HCs at either the cheek or antecubital fossa. In contrast, distinct species were associated with AD subgroups depending on the presence of co-occurring FA or FS (Fig. 2a; Table E3).

**Figure 2.**
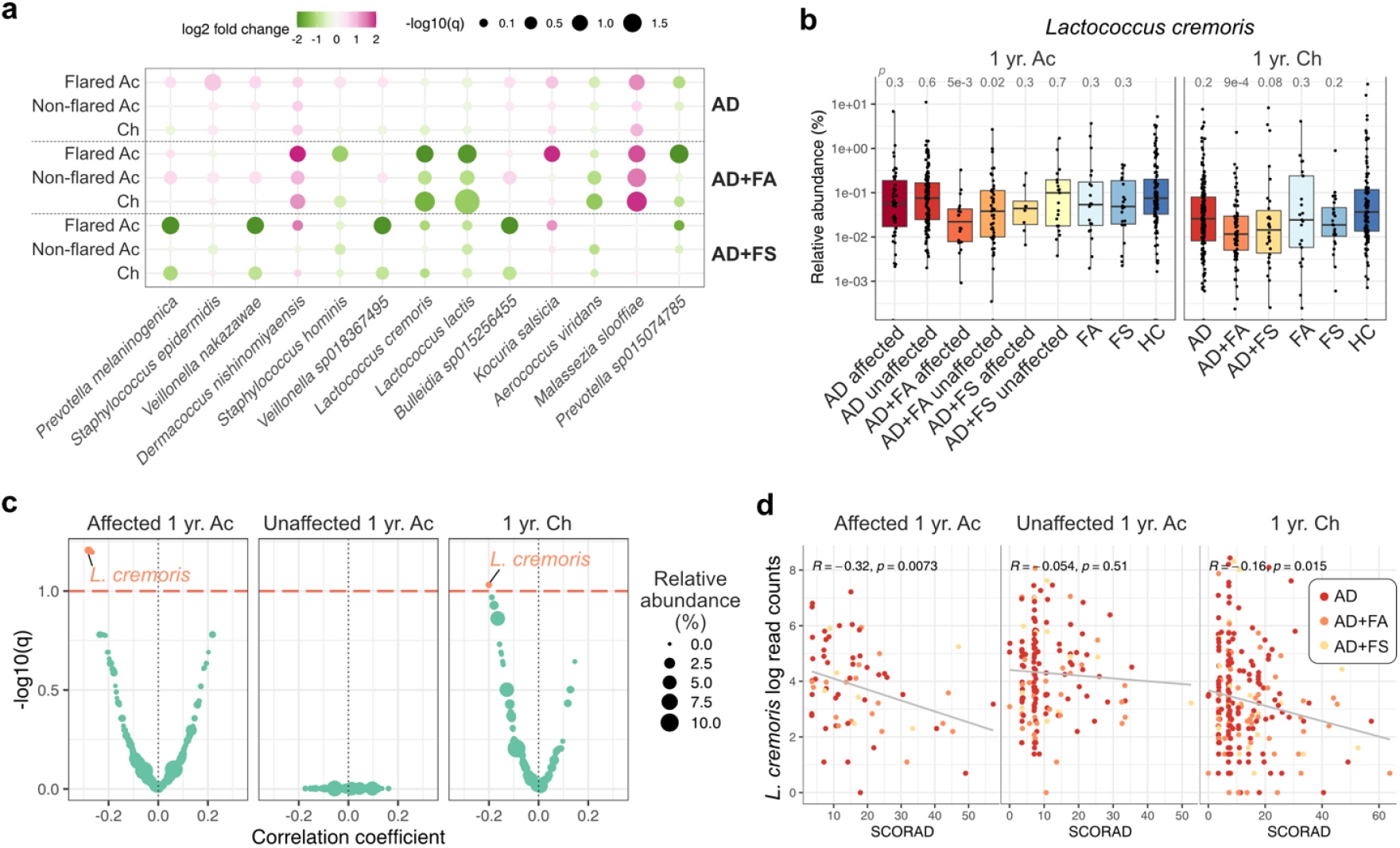
Microbial signatures associated with atopic disease. (**a**) Differentially abundant microbial taxa in the skin of infants with atopic disease compared to healthy controls. Taxa are arranged from left to right in descending order of abundance. (**b**) Relative abundance of *Lactococcus cremoris* in infants stratified by disease status and skin AD-affected status. Adjusted p-values from Mann–Whitney U tests comparing each group to healthy controls are displayed. (**c**) Pearson correlations between microbial abundance and SCORAD scores. P-values were adjusted using the Benjamini–Hochberg procedure. The horizontal dashed line indicates the significance threshold of q < 0.1; taxa with significant correlations are highlighted in orange. (**d**) Correlation between log-transformed read counts of *Lactococcus cremoris* and SCORAD score. Read counts were rarefied to a uniform sequencing depth prior to natural log transformation. Pearson correlation coefficients and corresponding p-values are shown. Ac: antecubital fossa. Ch: cheek. AD: atopic dermatitis. FA: food allergy. FS: food sensitization. HC: healthy control. SCORAD: scoring atopic dermatitis.

In infants with AD alone, *Staphylococcus epidermidis* was enriched in the AD-affected antecubital fossa compared to HCs, but not in unaffected skin or in infants with co-occurring FA or FS (Fig. E2a). More pronounced differences were observed in infants with both AD and FA, in whom *S. hominis* was decreased and *Dermococcus nishinomiyaensis* was enriched in the AD-affected antecubital fossa (Fig. E2b,c). Among fungi, *Malassezia slooffiae* was the only AD-associated species, showing increased abundance in infants with AD and FA regardless of affected status (Fig. E2d). Although *Staphylococcus aureus* has been linked to AD severity and flares, its abundance did not differ across AD subgroups in either affected or unaffected skin (Fig. E2e), consistent with the predominance of mild AD in this cohort.

Additionally, *Lactococcus cremoris* and *L. lactis* were reduced on both the cheeks and AD-affected antecubital fossa of infants with AD and FA compared to HCs (Fig. 2b). Notably, *L. cremoris* was the only taxon whose abundance negatively correlated with AD severity (SCORAD) across both sites (Fig. 2c,d). In infants with AD and FS, these shifts were absent; instead, *Prevotella melaninogenica* and *Veillonella* species were reduced in the AD-affected antecubital fossa (Fig. E2f). Together, these results indicate that infants with AD alone, AD with FA, and AD with FS are characterized by distinct microbial signatures.

### Altered skin microbiome at 2-3 months prior to atopic disease onset

Next, we investigated whether the skin microbiome of infants was already altered at 2-3 months, prior to the diagnosis of atopic diseases. At this timepoint, all infants were exclusively breastfed, and none had a parent-reported doctor diagnosis of AD, a clinical history of itchy skin, or prior use of topical steroids. Although no species were associated with the subsequent development of AD alone, five species were enriched on the skin of infants who developed both AD and FA by 1 year relative to HCs (Fig. 3a; Table E4), including *S. aureus* (Fig. 3b) and *Corynebacterium kefirresidentii* (Fig. E3a). In addition, *S. saprophyticus* was elevated on the antecubital fossa of infants who later developed both AD and FS (Fig. E3b). These findings indicate that early enrichment of specific bacterial species on the skin is associated with subsequent diagnoses of AD in combination with FA or FS, but not with AD alone.

**Figure 3.**
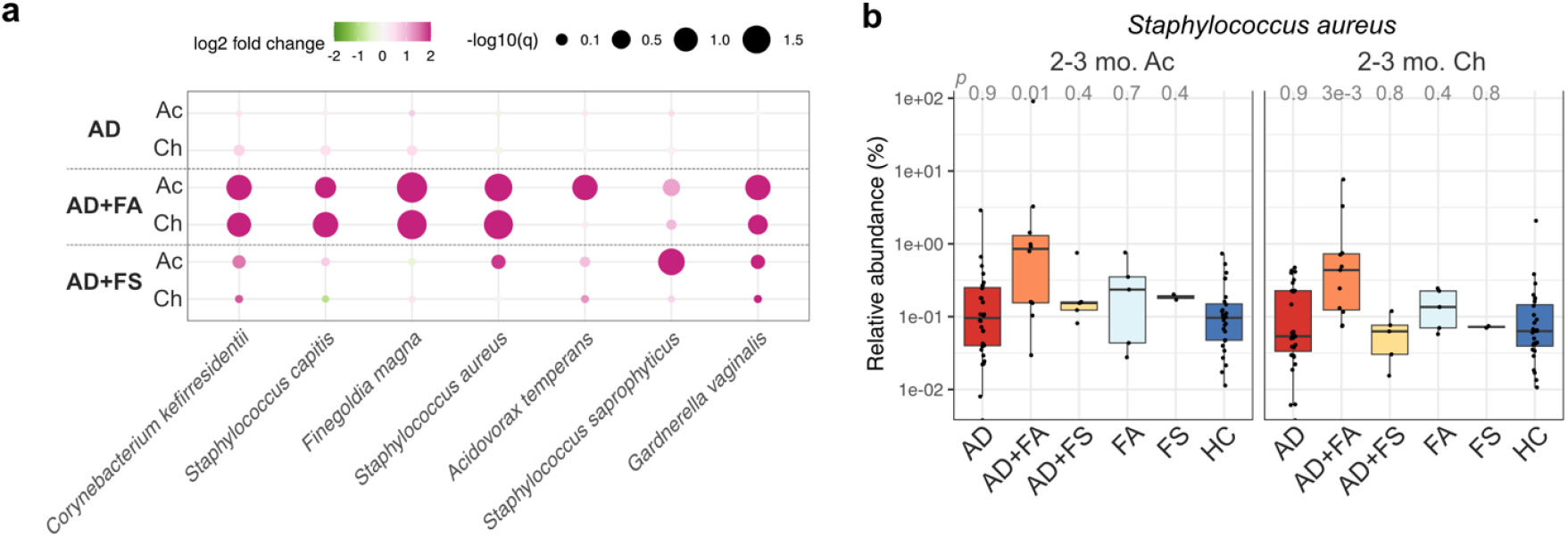
Skin microbial taxa at 2–3 months associated with atopic disease at 1 year. (**a**) Differentially abundant taxa in the skin of infants who developed atopic disease compared to healthy controls. Taxa are arranged from left to right in descending order of abundance. (**b**) Relative abundance of *Staphylococcus aureus* in infants stratified by disease status. Adjusted p-values from Mann–Whitney U tests comparing each group to healthy controls are displayed. Ac: antecubital fossa. Ch: cheek. AD: atopic dermatitis. FA: food allergy. FS: food sensitization. HC: healthy control.

### Impact of *FLG* mutations on infant skin microbiome in AD

Despite the strong associations of both host genetics and the skin microbiome with AD, the interaction between host genetics and the microbiome has been largely unexplored in the context of AD. In this study, we leveraged human DNA data captured in the shotgun sequencing datasets to investigate the interplay between host genetics and the skin microbiome. Given the reduced coverage of the human genome achieved by shotgun metagenomics compared to whole genome sequencing, we applied stringent criteria for calling genetic variants (Methods; Fig. E4a, b).

We focused our analysis on the *FLG* gene, which encodes the epidermal protein filaggrin that is crucial for maintaining the integrity of the skin barrier. Loss-of-function variants in *FLG* are the strongest genetic risk factor for AD. We identified 46 infants with AD and 14 without AD who possessed at least one *FLG* null mutation. The most frequent *FLG* null mutations in this infant cohort were R501X (i.e., Arg501*) and 2282del4 (i.e., Ser761fs) (Fig. 4a), which have been previously associated with AD in other cohorts^27–30^.

**Figure 4.**
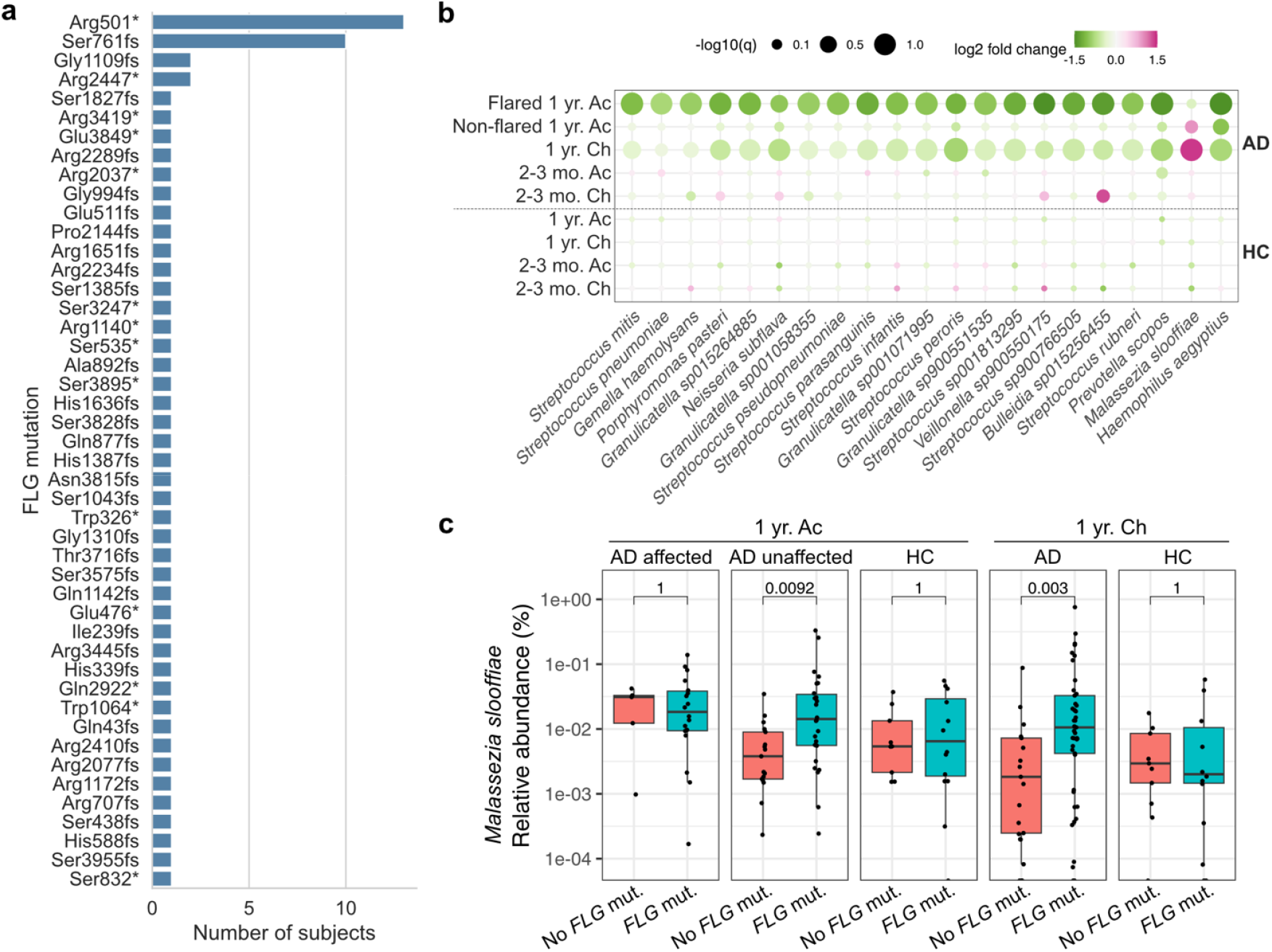
Association of *FLG* mutations with infant skin microbiome. (**a**) Summary of *FLG* null mutations identified in infants. (**b**) Differentially abundant skin taxa associated with *FLG* mutations, stratified by age, skin site, and AD status. Taxa are arranged from left to right in descending order of abundance. (**c**) Relative abundance of *Streptococcus mitis* in the skin of 1-year-old infants, stratified by disease status and by the presence or absence of *FLG* mutations. Adjusted p-values from Mann–Whitney U tests are shown. Ac: antecubital fossa. Ch: cheek. AD: atopic dermatitis. FA: food allergy. FS: food sensitization. HC: healthy control.

To identify taxa associated with host genetics, we compared the skin microbiome of infants with and without *FLG* mutations using differential abundance analysis. No taxa differed significantly between healthy infants with or without *FLG* mutations, nor were any microbial signatures associated with *FLG* mutations at 2-3 months (Fig. 4b; Table E5). In contrast, 17 species across 8 genera, most prominently *Streptococcus* and *Granulicatella*, were decreased on the AD-affected antecubital fossa of infants with *FLG* mutations compared to those without (Fig. 4b).

Some of these species were also reduced on the cheeks of infants with *FLG* mutations, including *Streptococcus peroris* (Fig. E4c). *Malassezia slooffiae* was the only taxon increased in infants with *FLG* mutations, but only on cheeks and unaffected antecubital fossa (Fig. 4c). None of these species showed differences at 2-3 months (Fig. E4d,e), suggesting that associations between *FLG* mutations and the skin microbiome may emerge only after clinical manifestations of allergic disease become apparent.

### Microbial functions associated with atopic diseases

To explore the microbial functions associated with atopic diseases, we applied a microbial GWAS (mGWAS) method to identify species-level, disease-associated microbial genetic variations using metagenomic data. We focused on five atopy-associated species (*S. epidermidis, S. aureus, D. nishinomiyaensis, L. lactis*, and *S. mitis*) and four highly abundant species in infants (*C*. acnes, *P. pasteri, M. luteus*, and *S. thermophilus*). We identified variants in *S. aureus, L. lactis*, and *S. thermophilus* that correlated significantly with SCORAD scores.

In *S. aureus*, we detected a missense mutation in the *fusA* gene that was significantly associated with lower SCORAD scores (Fig. 5a, b). In *S. thermophilus*, missense mutations in the *prs* gene correlated with higher SCORAD scores (Fig. 5c, d). In *L. lactis*, multiple genetic variants, including mutations in *ccd* and *ypiA*, were associated with AD severity (Fig. E5a–c). Taken together, these findings highlight strain-level species-specific adaptations that may shape microbial contributions to AD. Genetic variations affecting antibiotic resistance and metabolic pathways offer plausible mechanistic links between microbial function and disease severity, pointing to potential microbial targets for early-life intervention.

**Figure 5.**
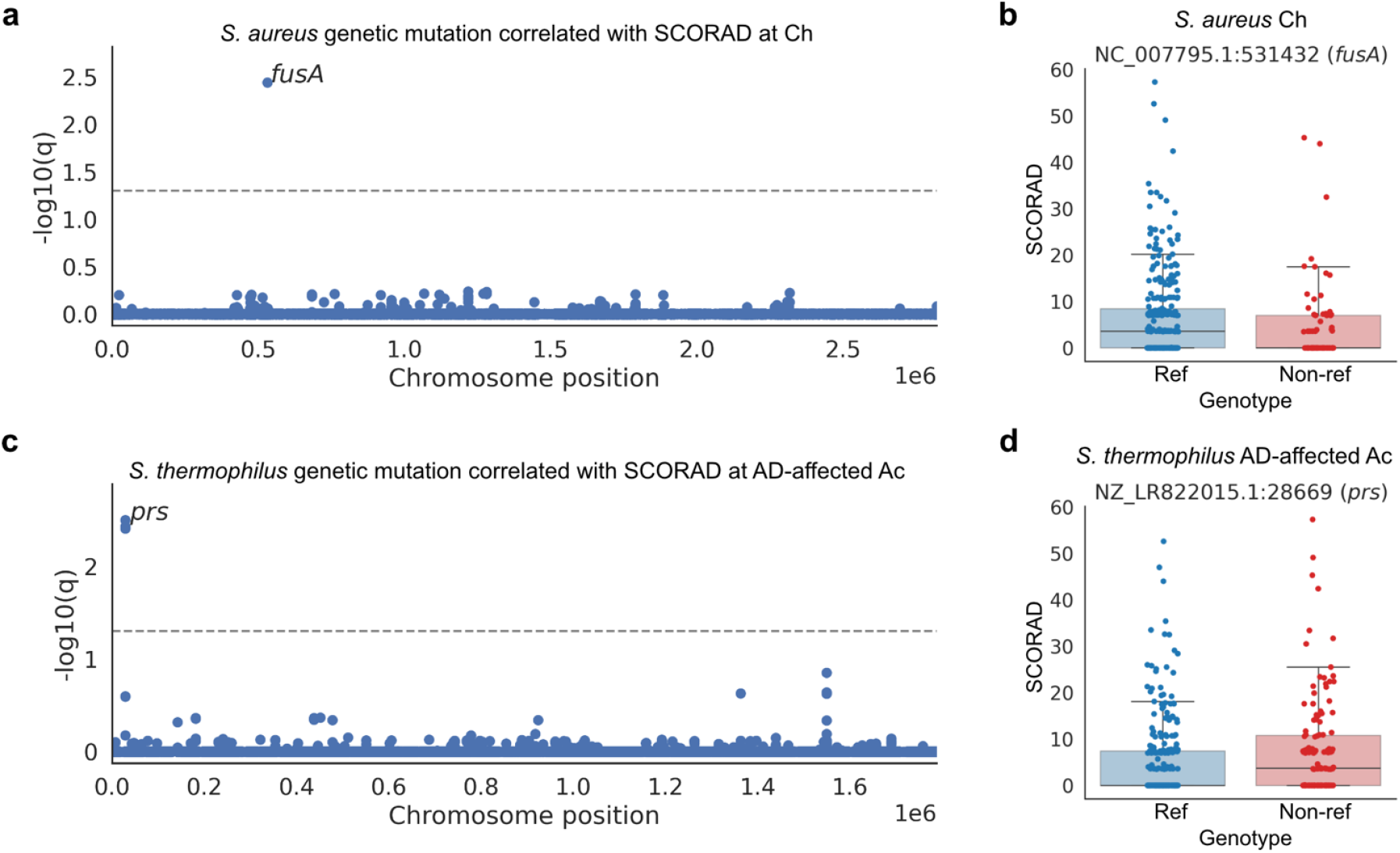
Microbial functions associated with atopic diseases identified by microbial GWAS. (**a**) Manhattan plot showing adjusted *p*-values for associations between *Staphylococcus aureus* variants and SCORAD scores in cheek samples. (**b**) Missense mutation in the *fusA* gene of *S. aureus* showing a negative association with SCORAD score. (**c**) Manhattan plot showing adjusted *p*-values for associations between *Streptococcus thermophilus* variants and SCORAD scores in AD-affected antecubital fossa samples. (**d**) Missense mutations in the *prs* gene of *S. thermophilus* showing a positive association with SCORAD score. Ac: antecubital fossa. Ch: cheek. SCORAD: scoring atopic dermatitis. Dashed line on Manhattan plot indicates the significance threshold of q < 0.05.

### Strain sharing of disease-associated skin microbiome between infants and mothers

Disease-associated microbial genetic variations and functions reveal strain-level specificity of the skin microbiome in relation to the infant’s clinical status. To explore the source of disease-associated strains, we predicted strain types from the metagenomic datasets of related infant-mother pairs using StrainGE^31^ for 6 species, including 5 species associated with atopic disease (*S. epidermidis, S. aureus, D. nishinomiyaensis, L. lactis*, and *L. cremoris*) plus *C. acnes*. Each species displayed varying numbers of predicted strains per individual (Fig. 6a). For instance, individuals typically harbored only one *S. aureus* strain but a median of three *S. epidermidis* strains. These findings align with prior work using saturated culturing from 20 subjects^13^.

**Figure 6.**
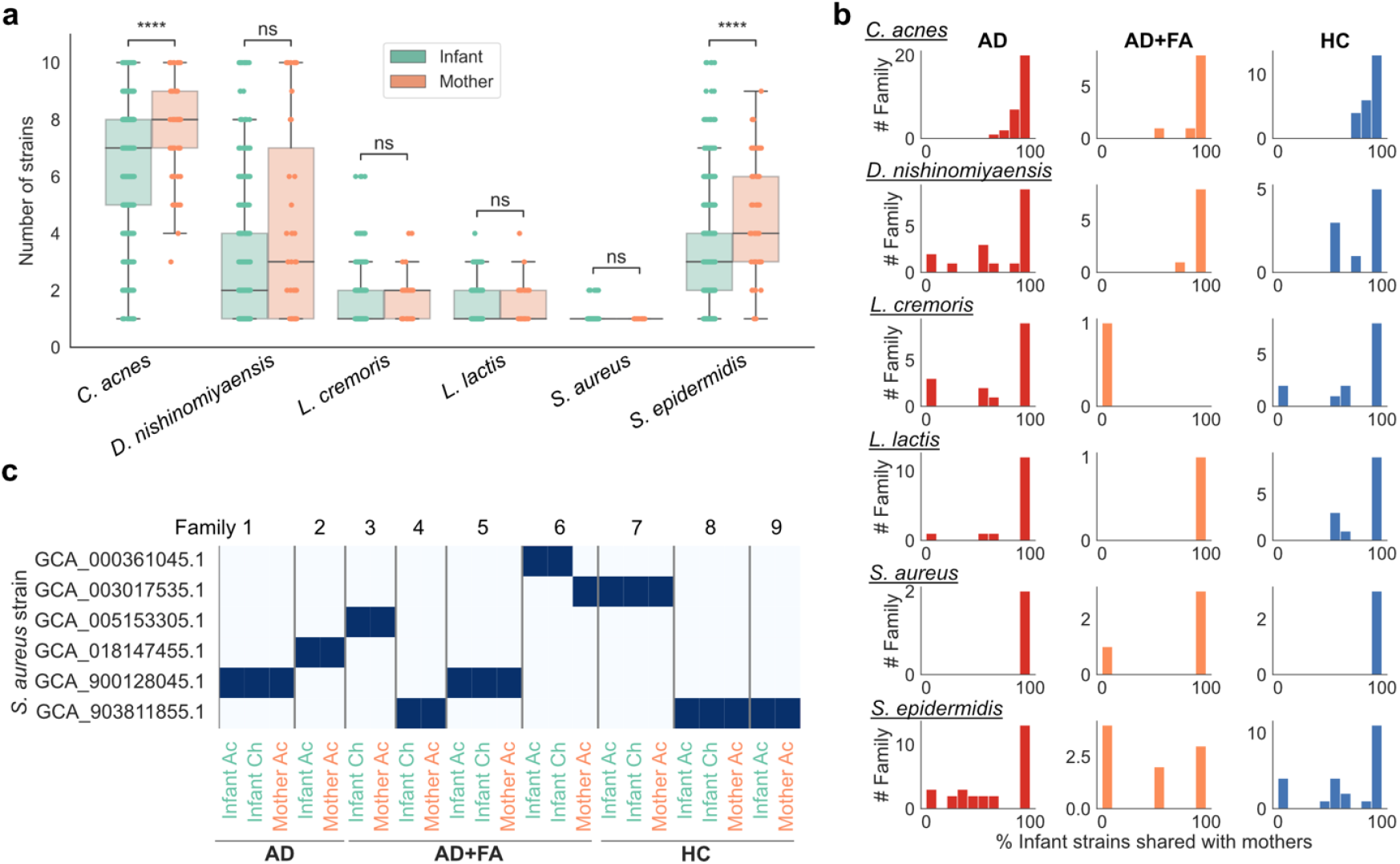
Familial strain sharing between infants and mothers. (**a**) Number of strains identified for atopic disease–associated species and *Cutibacterium acnes*. Comparisons between infants and mothers were performed using Mann–Whitney U tests. ****P < 0.0001. (**b**) Number of families in which strains found on infants are also detected on mothers, stratified by infant disease status. (**c**) Strain sharing of *Staphylococcus aureus* within families. Grey vertical lines separate families. Ac: antecubital fossa. Ch: cheek. AD: atopic dermatitis. FA: food allergy. FS: food sensitization. HC: healthy control.

Notably, species like *S. epidermidis* and *C. acnes* exhibited significantly greater strain diversity on mothers compared to infants, reflecting reduced strain diversity in the infant skin microbiome.

Comparing strain types between mothers and infants revealed a significant overlap across all tested species regardless of infant disease status (permutation test adjusted p-value < 0.01; Fig. 6b). For instance, *S. aureus* was detected in both infant and maternal skin in 9 families, with 8 of these dyads sharing the same *S. aureus* strain (Fig. 6c). Consistent with previous findings based on cultured isolates^26^, *C. acnes* also exhibited strain sharing between infants and mothers in our metagenomic analysis (Fig. 6b).

Certain species also exhibited significant temporal persistence of strains on infant skin across visits, including *C. acnes* and *S. aureus* (permutation test adjusted p-value < 0.01; Fig. E6a). For example, among six infants with detectable *S. aureus* at two time points, all retained at least one strain from 2-3 months to 1 year of age (Fig. E6b). Notably, two infants carried two distinct strains at 2-3 months, but one strain was lost by 1 year. In contrast, species such as *Staphylococcus epidermidis* and *Lactococcus lactis* showed a low proportion of persistent strains, suggesting that strains of these disease-associated species are more likely acquired later in infancy.

## DISCUSSION

While increasing attention has focused on infant gut dysbiosis in allergic disease, our study identifies the skin microbiome as an independent and early site of microbial divergence that precedes and stratifies allergic outcomes. Using deep shotgun metagenomic sequencing at unprecedented scale in infant skin, we identified microbial features that emerged before disease diagnosis and distinguished infants who developed AD alone from those who developed AD with co-occurring FA or FS. Unlike prior studies that largely contrasted AD versus healthy controls, our work resolves microbiome differences across clinically distinct AD phenotypes, providing insight into heterogeneity within AD. A further strength is the use of a population-based cohort with well-phenotyped outcomes, including FA confirmed by oral food challenge.

Within AD, overall microbial diversity was similar across subtypes, but we observed distinct taxonomic signatures associated with specific clinical combinations. *S. epidermidis* was enriched in infants with AD alone, while *S. hominis* was decreased in infants with both AD and FA. Both these species have previously been associated with AD^10,12^, but our results suggest that the association of staphylococci with AD may correlate with FA co-occurrence and may partially explain why some infants with AD also develop FA and others do not. Although several prior studies have associated *S. aureus* overgrowth with AD severity and flares, we did not observe differences in *S. aureus* abundance across AD subgroups at 12 months in either affected or unaffected skin. This may reflect the predominance of mild AD in our cohort, consistent with a less prominent role for *S. aureus* in milder disease^12,32^.

Beyond staphylococci, we identified additional microbes specific to the co-occurrence of AD and FA, including increased *Dermacoccus nishinomiyaensis* and decreased *Lactococcus* species. Although depletion of *D. nishinomiyaensis* has previously been linked to AD flares^9^, its enrichment in the skin of infants with AD and FA may reflect a compensatory response to restore immune homeostasis^33–35^. In contrast, the reduction of *Lactococcus*, which has been shown to ameliorate AD in murine models and a small clinical study^36^, suggests a protective role in disease pathogenesis. *Lactococcus* abundance was also inversely correlated with AD severity, highlighting its relevance for human disease and potential therapeutic utility. Moreover, because shotgun metagenomics profiles all kingdoms, our analysis also revealed a fungal signal: *Malassezia slooffiae* showed elevated abundance in infants with AD and FA. *Malassezia* spp. are known to elicit pro-inflammatory responses by interacting directly with the host immune system and by impairing skin barrier function. Notably, infants with severe AD and FA have previously been shown to have a higher likelihood of sensitization to *Malassezia* species^37^.

Infants with AD and FS exhibited a distinct microbial signature characterized by reduced *Prevotella* and *Veillonella* species. In the gut, depletion of these taxa has been associated with increased risk of AD, FS, and FA, potentially through reduced short-chain fatty acid production. Our findings suggest that their reduction on skin may similarly contribute to atopic risk^38–40^.

Notably, we identified early-life skin dysbiosis as being specifically associated with the later co-occurrence of AD with FA or FS, rather than serving as a universal precursor to AD. Infants who later developed AD with FA or FS showed increased early-life abundance of *Staphylococcus* species, with *S. aureus* specifically enriched in those who developed AD with FA. While prior studies have linked early-life skin dysbiosis to subsequent AD^15,16^ or AD with FA^41^, our findings refine these associations by showing that early skin dysbiosis is evident only in infants who later develop AD with allergic co-occurrence, and not in those who develop AD alone. These results further refine prior links between *S. aureus* colonization and the co-occurrence of AD and FA in children^18,19^ by showing that *S. aureus* enrichment occurs before clinical disease onset.

*FLG* null mutations are a major genetic risk factor for AD, and their relationship with skin dysbiosis has been examined in prior studies^42–45^. Here, we present a distinct analytical framework that infers host *FLG* genotypes directly from skin metagenomic data, enabling joint analysis of host genetics and the skin microbiome. Unlike prior studies focused primarily on *S. aureus* and *S. epidermidis*^42,44^, our approach extends *FLG*–microbiome associations across the broader skin microbial community. Using this framework, we identified reduced abundance of *Streptococcus* species and increased abundance of *Malassezia slooffiae* in *FLG* mutation carriers compared to non-carriers. Given the pro-inflammatory potential of certain *Streptococcus* species^19^, we speculate that, in individuals without a genetic predisposition to barrier dysfunction, microbial dysbiosis may play a more prominent role in driving skin inflammation. In contrast, impaired barrier function may preferentially support colonization by lipid-dependent fungi such as *Malassezia slooffiae*.

To move beyond taxonomic associations, we examined strain-level genetic variation using shotgun metagenomics. We identified a missense mutation in the *fusA* gene of *S. aureus* associated with lower AD severity. Given the known links between *fusA* mutations and fusidic acid resistance^46^, this association may reflect prior antibiotic exposure or other selection pressures rather than a direct effect on disease severity. In *S. thermophilus*, a mutation in the *prs* gene implicated altered nucleotide biosynthesis, correlating with higher AD severity, raising the possibility that altered nucleotide biosynthesis pathways in this commensal species might influence skin inflammation or host–microbe interactions. We acknowledge that most disease-associated genes and pathways still require further validation. Despite this, our findings support the presence of disease-associated strain-level variation in the skin microbiome, pointing to an important future direction for microbiome research.

While prior work has demonstrated mother-infant strain sharing of commensal skin microbiota in early life^26^, our study extends these observations by showing that skin microbial species associated with AD also exhibit strain sharing between infants and their mothers in the context of atopic disease. This suggests that early-life transmission of skin-associated microbes may contribute to the establishment of AD-associated skin microbial communities, potentially influencing disease risk beyond host genetics alone. Future studies that directly trace the origin and transmission routes of AD-associated skin strains will be essential for clarifying microbial sources and informing early-life preventive strategies.

This study has several limitations that should be considered. First, the cohort was entirely born in Australia, and generalizability to other populations requires caution. Additionally, the sampling point for disease onset was set at one year, when most cases had been diagnosed, but we cannot exclude cases classified as healthy may develop AD or FA later. We also cannot exclude the possibility that some infants may have had subclinical disease at 2–3 months. Moreover, most AD-affected infants in this study had mild to moderate symptoms, so caution is needed when extending the conclusions of this study to infants with severe AD. Finally, sample sizes for FA-only and FS groups, as well as for early-life samples (2–3 months), were limited, constraining statistical power for these analyses; therefore, null findings in these groups should not be interpreted as biological absence of effect.

Collectively, our study advances the field by extending early-life skin dysbiosis associations beyond AD to encompass allergic co-occurrence, host genetics, and microbial functional variation, providing a refined framework for understanding atopic disease heterogeneity and for identifying infants at risk for allergic comorbidities.

## ACKNOWLEDGMENTS

ZS is supported by the NIH Pathway to Independence Award R00 (AR084058). This research was supported by the National Institutes of Health (NIH) Intramural Research Program, including the National Institute of Allergy and Infectious Diseases (NIAID) and the National Human Genome Research Institute (NHGRI). Research reported in this publication was performed in part as a project of the Immune Tolerance Network and supported by the National Institute of Allergy and Infectious Diseases (NIAID) of the National Institutes of Health (NIH) under Award Number UM1AI109565. The contributions of the NIH author(s) are considered Works of the United States Government. The findings and conclusions presented in this paper are those of the author(s) and do not necessarily reflect the views of the NIH or the U.S. Department of Health and Human Services. The Vitality study was funded by the National Health and Medical Research Council (NHMRC) Australia (APP1146913); the National Institute of Health (NIH) Immune Tolerance Network; the DHB foundation; the Rotary Club of Camberwell; the Isabel and John Gilberston Charitable Trust; the Ilhan Food Allergy Foundation; the Kimberly Foundation Australia; the Constantinou Foundation; and other Philanthropic donations. Research at MCRI is supported by the Victorian Government’s Operational Infrastructure Support Program. KPP is supported by a NHMRC fellowship (GNT2008911) and a Melbourne Children’s Clinician-Scientist Fellowship. Audrey Walsh is supported by an Australian Government Research Training Program Scholarship provided by the Australian Commonwealth Government and the University of Melbourne, and a PhD scholarship from the Australian Government funded National Allergy Centre of Excellence (NACE), hosted by the Murdoch Children’s Research Institute (MCRI).

Disclosure of potential conflict of interest: KPP’s institution has received research grants from Aravax, DBV Technologies, Novartis and Siolta and consultant fees from Aravax, Novartis, RAPT Therapeutics, outside the submitted work. KPP is a non-executive director of OmnisOva.

We acknowledge the computational resources provided by the NIH HPC Biowulf Cluster (http://hpc.nih.gov). We thank Dr. Sean Conlan for feedback on the manuscript and assistance with data upload. We are grateful to the VITALITY families for their participation. OpenAI’s ChatGPT was used to assist with improving grammar, clarity, and readability; all scientific content, analyses, and interpretations remain the sole responsibility of the authors.

## REFERENCES

1. Loh W, Tang MLK. The Epidemiology of Food Allergy in the Global Context. Int J Environ Res Public Health. 2018;15:2043.

2. Langan SM, Irvine AD, Weidinger S. Atopic dermatitis. The Lancet. 2020;396:345–60.

3. Hadi HA, Tarmizi AI, Khalid KA, Gajdács M, Aslam A, Jamshed S. The Epidemiology and Global Burden of Atopic Dermatitis: A Narrative Review. Life. 2021;11:936.

4. Napolitano M, Fabbrocini G, Martora F, Genco L, Noto M, Patruno C. Children atopic dermatitis: Diagnosis, mimics, overlaps, and therapeutic implication. Dermatol Ther [Internet]. 2022 [cited 2025 Oct 5];35. Available from: https://onlinelibrary.wiley.com/doi/10.1111/dth.15901

5. Christensen MO, Barakji YA, Loft N, Khatib CM, Egeberg A, Thomsen SF, et al. Prevalence of and association between atopic dermatitis and food sensitivity, food allergy and challenge□proven food allergy: A systematic review and meta□analysis. J Eur Acad Dermatol Venereol. 2023;37:984–1003.

6. Arehart CH, Daya M, Campbell M, Boorgula MP, Rafaels N, Chavan S, et al. Polygenic prediction of atopic dermatitis improves with atopic training and filaggrin factors. J Allergy Clin Immunol. 2022;149:145–55.

7. Budu-Aggrey A, Kilanowski A, Sobczyk MK, 23 and Me Research Team, Shringarpure SS, Mitchell R, et al. European and multi-ancestry genome-wide association meta-analysis of atopic dermatitis highlights importance of systemic immune regulation. Nat Commun. 2023;14:6172.

8. Marenholz I, Grosche S, Kalb B, Rüschendorf F, Blümchen K, Schlags R, et al. Genome-wide association study identifies the SERPINB gene cluster as a susceptibility locus for food allergy. Nat Commun. 2017;8:1056.

9. Chng KR, Tay ASL, Li C, Ng AHQ, Wang J, Suri BK, et al. Whole metagenome profiling reveals skin microbiome-dependent susceptibility to atopic dermatitis flare. Nat Microbiol. 2016;1:16106.

10. Kong HH, Oh J, Deming C, Conlan S, Grice EA, Beatson MA, et al. Temporal shifts in the skin microbiome associated with disease flares and treatment in children with atopic dermatitis. Genome Res. 2012;22:850–9.

11. Nakatsuji T, Chen TH, Narala S, Chun KA, Two AM, Yun T, et al. Antimicrobials from human skin commensal bacteria protect against Staphylococcus aureus and are deficient in atopic dermatitis. Sci Transl Med. 2017;9:eaah4680.

12. Byrd AL, Deming C, Cassidy SKB, Harrison OJ, Ng WI, Conlan S, et al. Staphylococcus aureus and Staphylococcus epidermidis strain diversity underlying pediatric atopic dermatitis. Sci Transl Med. 2017;9:eaal4651.

13. Saheb Kashaf S, Harkins CP, Deming C, Joglekar P, Conlan S, Holmes CJ, et al. Staphylococcal diversity in atopic dermatitis from an individual to a global scale. Cell Host Microbe. 2023;31:578–592.e6.

14. Severn MM, Williams MR, Shahbandi A, Bunch ZL, Lyon LM, Nguyen A, et al. The Ubiquitous Human Skin Commensal Staphylococcus hominis Protects against Opportunistic Pathogens. Torres VJ, editor. mBio. 2022;13:e00930–22.

15. Kennedy EA, Connolly J, Hourihane JO, Fallon PG, McLean WHI, Murray D, et al. Skin microbiome before development of atopic dermatitis: Early colonization with commensal staphylococci at 2 months is associated with a lower risk of atopic dermatitis at 1 year. J Allergy Clin Immunol. 2017;139:166–72.

16. Rapin A, Rehbinder EM, Macowan M, Pattaroni C, Lødrup Carlsen KC, Harris NL, et al. The skin microbiome in the first year of life and its association with atopic dermatitis. Allergy. 2023;78:1949–63.

17. Łoś-Rycharska E, Gołębiewski M, Sikora M, Grzybowski T, Gorzkiewicz M, Popielarz M, et al. A Combined Analysis of Gut and Skin Microbiota in Infants with Food Allergy and Atopic Dermatitis: A Pilot Study. Nutrients. 2021;13:1682.

18. Leung DYM, Calatroni A, Zaramela LS, LeBeau PK, Dyjack N, Brar K, et al. The nonlesional skin surface distinguishes atopic dermatitis with food allergy as a unique endotype. Sci Transl Med. 2019;11:eaav2685.

19. Tsilochristou O, Du Toit G, Sayre PH, Roberts G, Lawson K, Sever ML, et al. Association of Staphylococcus aureus colonization with food allergy occurs independently of eczema severity. J Allergy Clin Immunol. 2019;144:494–503.

20. Allen KJ, Panjari M, Koplin JJ, Ponsonby AL, Vuillermin P, Gurrin LC, et al. VITALITY trial: protocol for a randomised controlled trial to establish the role of postnatal vitamin D supplementation in infant immune health. BMJ Open. 2015;5:e009377.

21. Williams HC, Jburney PG, Pembroke AC, Hay RJ, Atopic Dermatitis Diagnostic Criteria Working Party. The U.K. Working Party’s Diagnostic Criteria for Atopic Dermatitis. III. Independent hospital validation. Br J Dermatol. 1994;131:406–16.

22. Zannino D. VITALITY Statistical Analysis Plan V1.0 9th September 2025 [Internet]. Murdoch Childrens Research Institute; 2025 [cited 2025 Oct 20] p. 2166931 Bytes. Available from: https://mcri.figshare.com/articles/report/VITALITY_Statistical_Analysis_Plan_V1_0_9th_September_2025/30185827/1

23. Tirosh O, Conlan S, Deming C, Lee-Lin SQ, Huang X, NISC Comparative Sequencing Program, et al. Expanded skin virome in DOCK8-deficient patients. Nat Med. 2018;24:1815–21.

24. Che Y, Han J, Harkins CP, Hou P, Conlan S, Deming C, et al. Restoration of the human skin microbiome following immune recovery after hematopoietic stem cell transplantation. Cell Host Microbe. 2025;33:1412–1427.e5.

25. Saheb Kashaf S, Proctor DM, Deming C, Saary P, Hölzer M, NISC Comparative Sequencing Program, et al. Integrating cultivation and metagenomics for a multi-kingdom view of skin microbiome diversity and functions. Nat Microbiol. 2021;7:169–79.

26. Shen Z, Robert L, Stolpman M, Che Y, Allen KJ, Saffery R, et al. A genome catalog of the early-life human skin microbiome. Genome Biol. 2023;24:252.

27. Gao PS, Rafaels NM, Hand T, Murray T, Boguniewicz M, Hata T, et al. Filaggrin mutations that confer risk of atopic dermatitis confer greater risk for eczema herpeticum. J Allergy Clin Immunol. 2009;124:507–513.e7.

28. González-Tarancón R, Sanmartín R, Lorente F, Salvador□Rupérez E, Hernández-Martín A, Rello L, et al. Prevalence of FLG loss□of□function mutations R501X, 2282del4, and R2447X in Spanish children with atopic dermatitis. Pediatr Dermatol. 2020;37:98–102.

29. Greisenegger E, Novak N, Maintz L, Bieber T, Zimprich F, Haubenberger D, et al. Analysis of four prevalent filaggrin mutations (R501X, 2282del4, R2447X and S3247X) in Austrian and German patients with atopic dermatitis. J Eur Acad Dermatol Venereol. 2010;24:607–10.

30. Palmer CNA, Irvine AD, Terron-Kwiatkowski A, Zhao Y, Liao H, Lee SP, et al. Common loss-of-function variants of the epidermal barrier protein filaggrin are a major predisposing factor for atopic dermatitis. Nat Genet. 2006;38:441–6.

31. Van Dijk LR, Walker BJ, Straub TJ, Worby CJ, Grote A, Schreiber HL, et al. StrainGE: a toolkit to track and characterize low-abundance strains in complex microbial communities. Genome Biol. 2022;23:74.

32. Delanghe L, De Boeck I, Van Malderen J, Allonsius CN, Van Rillaer T, Bron PA, et al. Mild atopic dermatitis is characterized by increase in non-staphylococcus pathobionts and loss of specific species. Sci Rep. 2024;14:23659.

33. Sowani H, Kulkarni M, Zinjarde S. Harnessing the catabolic versatility of Gordonia species for detoxifying pollutants. Biotechnol Adv. 2019;37:382–402.

34. Yadav M, Chaudhary PP, D’Souza BN, Ratley G, Spathies J, Ganesan S, et al. Diisocyanates influence models of atopic dermatitis through direct activation of TRPA1. Xu SZ, editor. PLOS ONE. 2023;18:e0282569.

35. Zeldin J, Chaudhary PP, Spathies J, Yadav M, D’Souza BN, Alishahedani ME, et al. Exposure to isocyanates predicts atopic dermatitis prevalence and disrupts therapeutic pathways in commensal bacteria. Sci Adv. 2023;9:eade8898.

36. Suzuki T, Nishiyama K, Kawata K, Sugimoto K, Isome M, Suzuki S, et al. Effect of the Lactococcus Lactis 11/19-B1 Strain on Atopic Dermatitis in a Clinical Test and Mouse Model. Nutrients. 2020;12:763.

37. Nowicka D, Nawrot U. Contribution of Malassezia spp. to the development of atopic dermatitis. Mycoses. 2019;62:588–96.

38. Chen C, Chen K, Kong M, Chang H, Huang J. Alterations in the gut microbiotas of children with food sensitization in early life. Pediatr Allergy Immunol. 2016;27:254–62.

39. Fan X, Zang T, Dai J, Wu N, Hope C, Bai J, et al. The associations of maternal and children’s gut microbiota with the development of atopic dermatitis for children aged 2 years. Front Immunol. 2022;13:1038876.

40. Mahdavinia M, Rasmussen HE, Botha M, Binh Tran TD, Van Den Berg JP, Sodergren E, et al. Effects of diet on the childhood gut microbiome and its implications for atopic dermatitis. J Allergy Clin Immunol. 2019;143:1636–1637.e5.

41. Aoyama R, Nakagawa S, Ichikawa Y, Inohara N, Yamazaki Y, Ito T, et al. Neonatal skin dysbiosis to infantile atopic dermatitis: Mitigating effects of skin care. Allergy. 2024;79:1618–22.

42. Van Mierlo MMF, Pardo LM, Fieten KB, Van Den Broek TJ, Schuren FHJ, Van Geel M, et al. The Skin and Nose Microbiome and Its Association with Filaggrin Gene Mutations in Pediatric Atopic Dermatitis. Dermatology. 2022;238:928–38.

43. Clausen ML, Agner T, Lilje B, Edslev SM, Johannesen TB, Andersen PS. Association of Disease Severity With Skin Microbiome and Filaggrin Gene Mutations in Adult Atopic Dermatitis. JAMA Dermatol. 2018;154:293.

44. Nath S, Kumari N, Bandyopadhyay D, Sinha N, Majumder PP, Mitra R, et al. Dysbiotic Lesional Microbiome With Filaggrin Missense Variants Associate With Atopic Dermatitis in India. Front Cell Infect Microbiol. 2020;10:570423.

45. Stamatas GN, Insel R, Sørensen N, Palleja A, Moll JM, Oddos T, et al. Shifts in Infant Skin Microbiome at 2 Months after Short-Term Emollient Use from Birth Are Associated with Reduced Prevalence of Atopic Dermatitis at 12 Months in a High-Risk Cohort. J Invest Dermatol. 2025;145:2640–2643.e11.

46. Chen HJ, Hung WC, Tseng SP, Tsai JC, Hsueh PR, Teng LJ. Fusidic Acid Resistance Determinants in Staphylococcus aureus Clinical Isolates. Antimicrob Agents Chemother. 2010;54:4985–91.

