## Supplemental Materials and Methods for "Skin microbiome composition and function in the development of atopic diseases during infancy"

**Supplementary Materials and Methods**

**Detailed study design and participant selection**

The VITALITY trial is a randomized, double-blind, placebo-controlled trial with the primary aim to determine whether oral vitamin D supplementation during the first year of life lowers the risk of food allergy at age 12 months. This cohort of healthy, term, predominantly breastfed, vitamin D un-supplemented infants was recruited at 6-12 weeks postnatal age through council-run 2-month vaccination sessions and maternal-child-health clinics and online including social media in metropolitan Melbourne, Australia. Diagnosis of food allergy was determined at the 12-month clinic visit. Infants in the FA group had confirmed IgE-mediated food allergy to at least one of the 10 most common food allergens (egg, peanut, cow’s milk, cashew, sesame, shellfish, almond, soybean, hazelnut, wheat) derived from the algorithm which incorporates data from the oral food challenge, skin prick testing and parent-reported allergen ingestion and reaction history^1^. Infants with FS had positive food-specific IgE testing (as measured by skin prick testing and/or ImmunoCAP) to at least one of the 10 protocol-specified food allergens but were tolerant of all 10 foods. Infants in the AD group were diagnosed by clinical examination at the 6-month and/or 12-month clinic visits, when a SCORAD was also completed to document AD severity^2^. Infants in the healthy control group had no AD, no FA, and no sensitization to the 10-protocol specified foods or house dust mite (defined by negative skin prick testing and ImmunoCAP). Written informed consent was obtained for all participants in this study.

This work was conducted as a nested, phenotype-enriched microbiome study within the prospective VITALITY trial. We analyzed 1,078 skin swabs from 429 infants, collected from the antecubital fossa and cheek at two time points: the baseline home visit (age 2–3 months; n = 80 infants) and the 12-month clinic visit (n = 425 infants). Infant inclusion was based on the availability of skin samples and prospectively ascertained clinical phenotypes. All available skin samples from the baseline visit were included without enrichment or exclusion, comprising 29 infants who later developed AD only, 11 with both AD and FA, 5 with both AD and FS, 5 with FA only, 2 with FS only, and 28 healthy controls (HCs). The smaller sample size at the baseline visit was primarily due to restrictions on in-person home visits during the COVID-19 pandemic. At the 12-month visit, sample selection was intentionally enriched for infants with atopic phenotypes to increase statistical power for microbiome comparisons across clinically defined groups. Specifically, we included the vast majority of available skin swabs from disease groups (≥85% in each group), including 169 infants with AD only, 72 with both AD and FA, 30 with both AD and FS, 22 with FA only, and 25 with FS only. A subset of 108 HCs was randomly selected from the pool of non-atopic infants with available skin swabs.

AD-affected status was assessed only for the antecubital fossa at 12 months, where samples were collected from affected sites in 43 infants with AD only, 19 with both AD and FA, and 8 with both AD and FS; AD-affected status of the cheek was not recorded. In addition, we analyzed 70 maternal skin samples from mothers of infants with AD only (n = 32), both AD and FA (n = 10), FS only (n = 1), or HCs (n = 27).

Because this sampling strategy intentionally enriches for disease phenotypes, the analyzed cohort does not reflect population-level disease prevalence. Accordingly, all analyses are designed to evaluate within-phenotype microbial differences conditional on clinical status, rather than to estimate disease incidence or risk.

**Sample collection and negative controls**
Puritan foam swabs were collected and stored in 100 µL of Yeast Cell Lysis Buffer (Lucigen) at −80°C and shipped on dry ice. Participants were asked to avoid bathing within 24 hours prior to sample collection. To explicitly control for environmental and reagent-associated contamination, air swabs were collected in parallel during each sampling session by exposing sterile swabs to the sampling environment without skin contact. Air swabs were processed alongside skin swabs through identical DNA extraction, library preparation, and sequencing workflows, ensuring comparable exposure to potential sources of contamination. These air swabs served as negative controls for downstream analyses.

**DNA extraction and metagenomic sequencing**

Library preparation, sequencing, and sample processing were performed as previously described^3–5^. Briefly, DNA libraries for Illumina sequencing were prepared using the Nextera XT DNA Library Preparation Kit (Illumina) according to the manufacturer’s instructions, with a modification to increase the AMPure XP bead clean-up volume from 30 µL to 50 µL. A total of 1 ng of extracted DNA was used as input for the fragmentation step, where DNA was simultaneously fragmented and tagged with sequencing adapters in a single-tube enzymatic reaction.

The prepared libraries were sequenced on the Illumina NovaSeqX+ platform at the NIH Intramural Sequencing Center, generating 2 × 150 bp reads and yielding a median of 90 million total reads per sample after quality control. Adapter sequences and low-quality bases were trimmed using Cutadapt v4.06^6^. To remove host-derived sequences, reads were aligned to the human reference genome (T2T-CHM13v2-0)^7^ using Bowtie v2.5.1^8^ , and mapped human reads were discarded.

Skin samples yielded a median of 96 million total reads per sample after quality control (IQR = 78-116 million), including 80 million human reads (IQR = 54-105 million) and 13 million non-human reads (IQR = 5-23 million).

Air swab controls were sequenced alongside skin swabs in the same batch, producing 100-fold fewer reads (IQR = 37-158 times fewer) compared to their corresponding skin swabs, which indicated minimal contamination. Four infant antecubital fossa samples and two infant cheek samples produced a similar or lower number of reads (<2-fold more) compared to their corresponding air controls and were excluded from further analysis. Four samples with less than 100,000 taxonomy-assigned reads were also excluded. This resulted in a final dataset comprising 998 infant and 70 maternal skin metagenomic samples for downstream analysis.

**Metagenomic data assembly and processing**

Shotgun metagenomic reads were assembled using an established pipeline developed for creating the Early-Life Skin Genome (ELSG) catalog^9^. The newly assembled MAGs were clustered with the ELSG catalog at a 95% ANI threshold using dRep v3.4.2 with the following parameters: “-pa 0.90 -sa 0.95 -nc 0.30 -cm larger --S_algorithm fastANI --multiround_primary_clustering --run_tertiary_clustering --clusterAlg single”^10^. Taxonomic annotations for prokaryotic MAGs were assigned using the “classify_wf” workflow in GTDB-Tk v2.1.0 with default settings, based on GTDB database release 207^11,12^. Metagenomic reads were mapped using Kraken v2.1.2 with the parameters “--confidence 0.1 --paired”^13^, and species-level microbial abundances were estimated with Bracken v2.5 using the parameters “-r 100 -l S”^14^. These analyses utilized the standard RefSeq database (release 211) as well as custom databases created from the SMGC and ELSG catalogs and the newly assembled MAGs, following an established protocol^9^. Two infant skin metagenomic datasets containing fewer than 100,000 classified microbial reads by Kraken were excluded from downstream analysis.

**Alpha and beta diversity**

Alpha diversity was assessed using the Shannon index, while beta diversity was quantified using Bray-Curtis dissimilarity. Associations between microbial diversity and clinical variables were assessed using PERMANOVA for beta diversity and ANOVA for alpha diversity. PERMANOVA analyses were conducted with the “adonis2” function in the vegan R package, using 9,999 permutations and “bray” method^15^. Age and skin site were tested across all samples, whereas the other variables were tested separately for each age group and skin site. P-values were corrected with the Benjamini–Hochberg procedure.

**Differential abundance analysis**

Differential abundance analysis was performed on raw classified reads using LinDA with the prevalence filter set to 0.3, the mean abundance filter set to 5e-5, and the Benjamini-Hochberg procedure applied for p-value adjustment^16^. Taxa with significant differential abundance were identified using a q-value cutoff of 0.1.

**Correlation analysis with AD severity**

### Rarefied reads were used for analysis. Taxa with a median relative abundance below 0.01% were filtered out to minimize the number of analyses. Pearson correlation coefficient and p-value were computed to examine the linear relationship between log-transformed abundances and SCORAD scores across all infants. Separate models were constructed for samples from AD-affected and unaffected skin, with both models including data from healthy controls. P-values were corrected with the Benjamini-Hochberg procedure.

***FLG* mutation identification**

Shotgun metagenomic reads were mapped to the hg38 genome using Bowtie v2.5.1 with default parameters^8^. Genetic variants in the *FLG* gene were identified with BCFtools v1.17 by specifying the target region as chr1:152302165-152325239 and using parameters “-m -P 1e-1”^17^. Variants were annotated for their predicted impacts using SnpEff v5.2^18^. Infants with at least one variant classified as having a “high” impact, supported by at least three reads, were categorized as having a *FLG* null mutation. Conversely, infants with no reads matching any of the high-impact variants identified in the cohort and at least three reads supporting the reference allele at each position were considered free of *FLG* null mutations.

Differential abundance analysis was conducted to compare microbial profiles between infants with and without *FLG* null mutations, using LinDA^16^. The prevalence filter was set at 0.3, and the mean abundance filter was increased to 1e-5 to reduce the number of tests due to the small sample size.

**Microbial GWAS**

Shotgun metagenomic reads were mapped to each reference genome with Bowtie v2.5.1 using “sensitive-local” mode^8^. Genetic variants were identified with GATK HaplotypeCaller using the parameters “--sample-ploidy 4 --native-pair-hmm-threads”^19^. Variants were annotated for impact using SnpEff v5.2^18^ based on custom-built databases, and those with moderate or high impact were selected for association tests. The presence or absence of variants was tested between two comparison groups using Fisher’s exact tests. Variants with adjusted p-values below 0.05 were considered statistically significant.

**Strain sharing analysis**

Complete genomes for the target species (*C. acnes, S. epidermidis, S. aureus, D. nishinomiyaensis, L. lactis,* and *L. cremoris*) were downloaded from the GenBank database (accessed October 11, 2024) and dereplicated using dRep v3.4.2 with the parameters “-pa 0.95 -sa 0.995 -nc 0.6 -cm larger --S_algorithm fastANI --multiround_primary_clustering --run_tertiary_clustering --clusterAlg single -comp 90 -con 5”^10^. The dereplicated genomes were then used to create species-specific databases with *straingst* tool of StrainGE v1.2^29^. The recommended pipeline included the following steps: “kmerize -k 31” for k-mer generation, “kmersim --all-vs-all -S jaccard -S subset” for k-mer similarity estimation, “cluster -d -C 2 -c 0.995” for clustering, and “createdb” to build the database. Metagenomic datasets were also sketched using the same k-mer size with the command “straingst kmerize -k 31.” Strain identification was conducted with the command “straingst run -i 10.”

Permutation tests were performed to evaluate the statistical significance of strain sharing between infants and their mothers. To generate a null distribution, identified strains from all individuals were randomly shuffled, and the median number of shared strains between related infant-mother pairs was recalculated. This process was repeated 1,000 times, producing a null distribution of median shared strains. The p-value was determined as the proportion of shuffled medians exceeding the observed median number of shared strains between infant-mother pairs. P-values < 0.01 were considered significant after the Benjamini–Hochberg correction.

**Data and code availability**

The raw metagenomic sequencing data are available in the NCBI BioProject database under project number PRJNA971252. The 250 new MAGs are available at <https://research.nhgri.nih.gov/projects/ELSG/downloads/InfantAD_mags_250v2.tar.zip>. Individual-level clinical and demographic data, as well as identified *FLG* mutations, may be accessed upon request. Applications to access data and/or samples can be submitted to or through the Melbourne Children’s Lifecourse Initiative (https://lifecourse.melbournechildrens.com/). All Python and R code for generating figures and the microbial GWAS pipeline are available on GitHub: <https://github.com/zeyang-shen/VITALITY_skin_atopy>.

**References**

1.	Zannino D. VITALITY Statistical Analysis Plan V1.0 9th September 2025 [Internet]. Murdoch Childrens Research Institute; 2025 [cited 2025 Oct 20] p. 2166931 Bytes. Available from: https://mcri.figshare.com/articles/report/VITALITY_Statistical_Analysis_Plan_V1_0_9th_September_2025/30185827/1

2.	Williams HC, Jburney PG, Pembroke AC, Hay RJ, Atopic Dermatitis Diagnostic Criteria Working Party. The U.K. Working Party’s Diagnostic Criteria for Atopic Dermatitis. III. Independent hospital validation. Br J Dermatol. 1994;131:406–16.

3.	Byrd AL, Deming C, Cassidy SKB, Harrison OJ, Ng WI, Conlan S, et al. *Staphylococcus aureus* and *Staphylococcus epidermidis* strain diversity underlying pediatric atopic dermatitis. Sci Transl Med. 2017;9:eaal4651.

4.	Tirosh O, Conlan S, Deming C, Lee-Lin SQ, Huang X, NISC Comparative Sequencing Program, et al. Expanded skin virome in DOCK8-deficient patients. Nat Med. 2018;24:1815–21.

5.	Che Y, Han J, Harkins CP, Hou P, Conlan S, Deming C, et al. Restoration of the human skin microbiome following immune recovery after hematopoietic stem cell transplantation. Cell Host Microbe. 2025;33:1412-1427.e5.

6.	Martin M. Cutadapt removes adapter sequences from high-throughput sequencing reads. EMBnetjournal Vol 17 No 1 Gener Seq Data Anal - 1014806ej171200 [Internet]. 2011; Available from: https://journal.embnet.org/index.php/embnetjournal/article/view/200

7.	Nurk S, Koren S, Rhie A, Rautiainen M, Bzikadze AV, Mikheenko A, et al. The complete sequence of a human genome. Science. 2022;376:44–53.

8.	Langmead B, Salzberg SL. Fast gapped-read alignment with Bowtie 2. Nat Methods. 2012;9:357–9.

9.	Shen Z, Robert L, Stolpman M, Che Y, Allen KJ, Saffery R, et al. A genome catalog of the early-life human skin microbiome. Genome Biol. 2023;24:252.

10.	Olm MR, Brown CT, Brooks B, Banfield JF. dRep: a tool for fast and accurate genomic comparisons that enables improved genome recovery from metagenomes through de-replication. ISME J. 2017;11:2864–8.

11.	Chaumeil PA, Mussig AJ, Hugenholtz P, Parks DH. GTDB-Tk v2: memory friendly classification with the genome taxonomy database. Borgwardt K, editor. Bioinformatics. 2022;38:5315–6.

12.	Parks DH, Chuvochina M, Chaumeil PA, Rinke C, Mussig AJ, Hugenholtz P. A complete domain-to-species taxonomy for Bacteria and Archaea. Nat Biotechnol. 2020;38:1079–86.

13.	Wood DE, Lu J, Langmead B. Improved metagenomic analysis with Kraken 2. Genome Biol. 2019;20:257.

14.	Lu J, Breitwieser FP, Thielen P, Salzberg SL. Bracken: estimating species abundance in metagenomics data. PeerJ Comput Sci. 2017;3:e104.

15.	Dixon P. VEGAN, a package of R functions for community ecology. J Veg Sci. 2003;14:927–30.

16.	Zhou H, He K, Chen J, Zhang X. LinDA: linear models for differential abundance analysis of microbiome compositional data. Genome Biol. 2022;23:95.

17.	Danecek P, Bonfield JK, Liddle J, Marshall J, Ohan V, Pollard MO, et al. Twelve years of SAMtools and BCFtools. GigaScience. 2021;10:giab008.

18.	Cingolani P, Platts A, Wang LL, Coon M, Nguyen T, Wang L, et al. A program for annotating and predicting the effects of single nucleotide polymorphisms, SnpEff: SNPs in the genome of Drosophila melanogaster strain w^1118^ ; iso-2; iso-3. Fly (Austin). 2012;6:80–92.

19.	McKenna A, Hanna M, Banks E, Sivachenko A, Cibulskis K, Kernytsky A, et al. The Genome Analysis Toolkit: A MapReduce framework for analyzing next-generation DNA sequencing data. Genome Res. 2010;20:1297–303.
