## Supplemental Figures for "Skin microbiome composition and function in the development of atopic diseases during infancy"

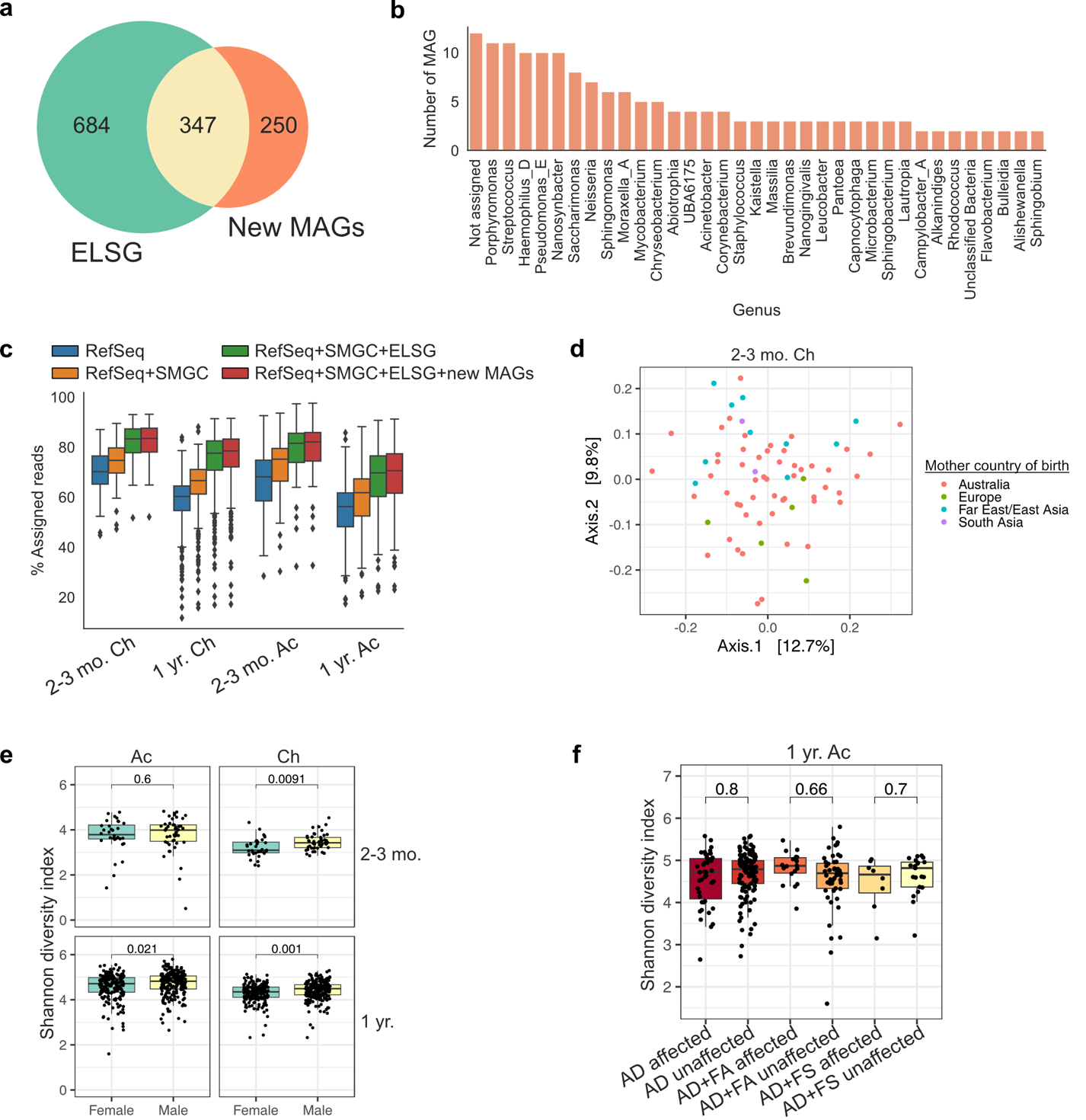

**Figure E1.** (**a**) Comparison of the ELSG catalog and new MAGs assembled from additional skin metagenomes. The Venn diagram shows the number of shared and unique MAGs. (**b**) Genus-level taxonomy of 250 new MAGs unique to the additional metagenomes. (**c**) Read classification rates using different reference databases. (**d**) Principal coordinate analysis of cheek metagenomes at 2–3 months of age, colored by mothers’ region of birth (proxy for ethnicity). (**e**) Shannon diversity index comparing male and female infants. (**f**) Shannon diversity index of AD-affected vs. unaffected skin in infants across disease groups. Adjusted p-values from Mann–Whitney U tests are shown. Ac: antecubital fossa. Ch: cheek. AD: atopic dermatitis. FA: food allergy. FS: food sensitization. HC: healthy control. SMGC: Skin Microbial Genome Collection. ELSG: Early-Life Skin Genome. MAG: metagenome-assembled genome.

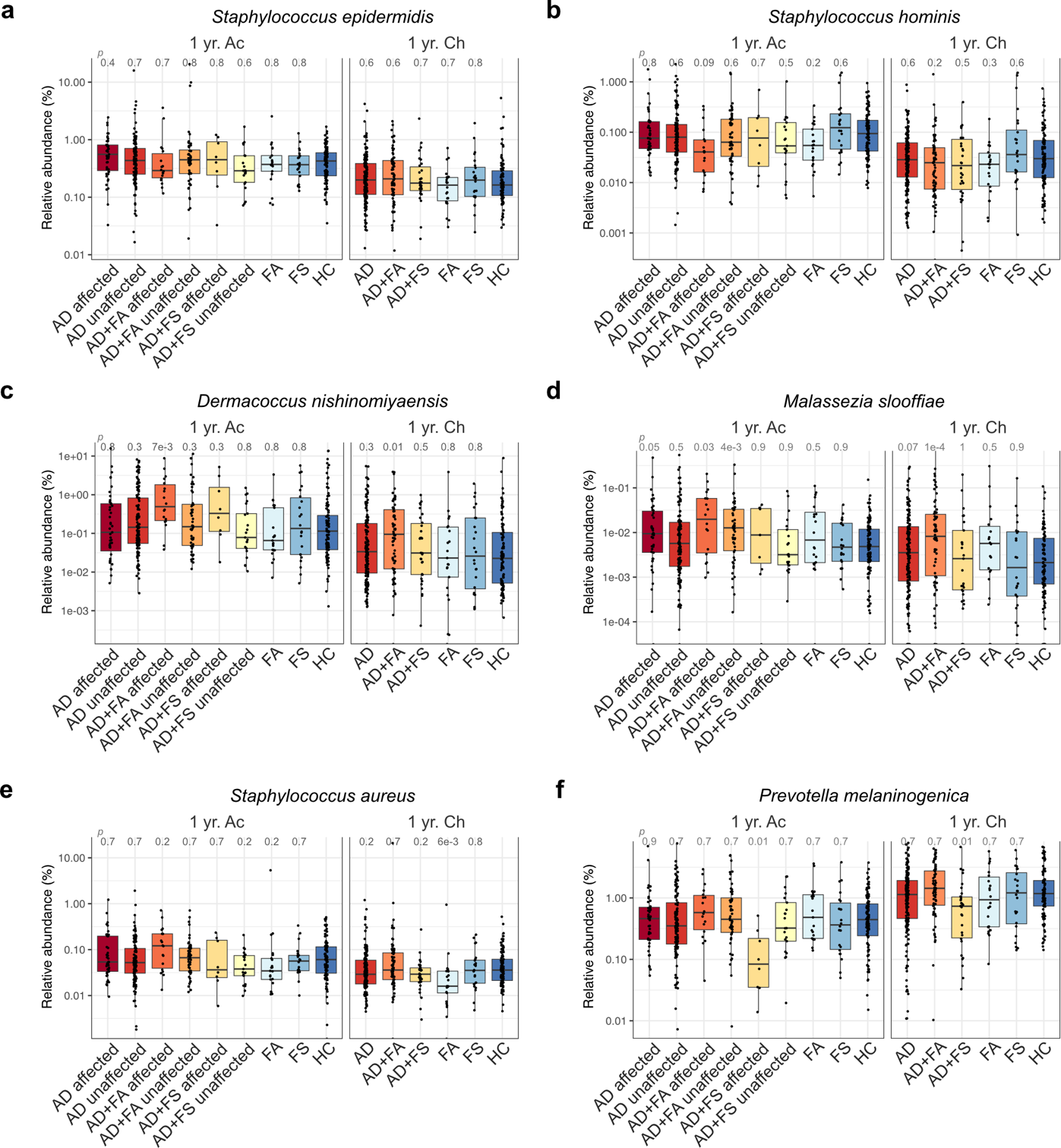

**Figure E2.** Relative abundance of (**a**) Staphylococcus epidermidis, (**b**) Staphylococcus hominis, (**c**) Malassezia slooffiae, (**d**) Dermacoccus nishinomiyaensis, (**e**) Prevotella melaninogenica, and (**f**) Staphylococcus aureus in infants stratified by disease status and AD-affected status. Adjusted p-values from Mann–Whitney U tests comparing each group to healthy controls are displayed. Ac: antecubital fossa. Ch: cheek. AD: atopic dermatitis. FA: food allergy. FS: food sensitization. HC: healthy control.

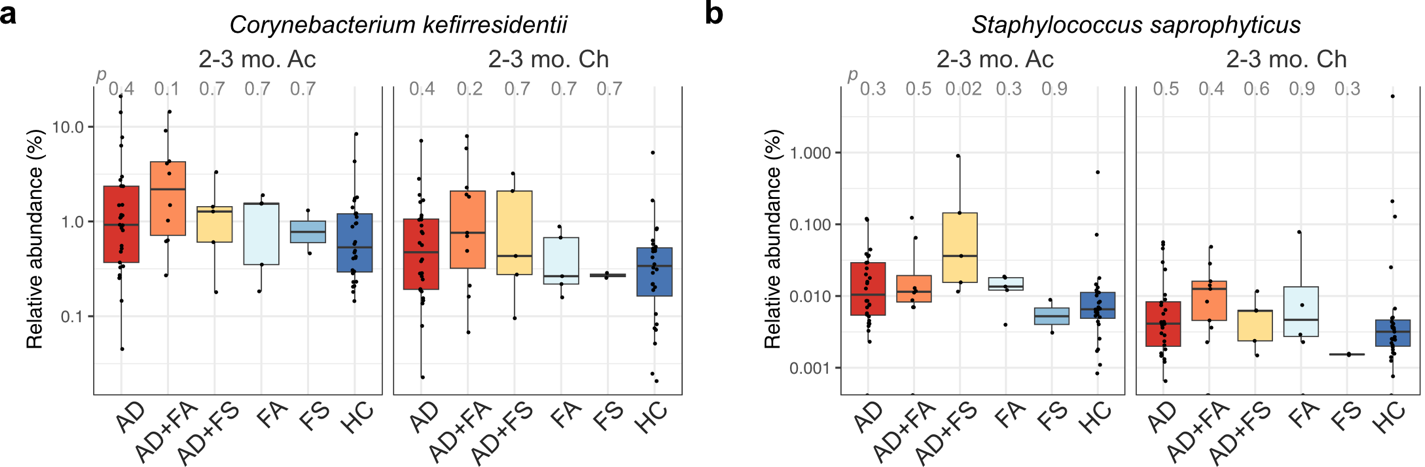

**Figure E3.** Relative abundance of (**a**) Corynebacterium kefirresidentii and (**b**) Staphylococcus saprophyticus in infants at 3 months stratified by disease status. Adjusted p-values from Mann–Whitney U tests comparing each group to healthy controls are displayed. Ac: antecubital fossa. Ch: cheek. AD: atopic dermatitis. FA: food allergy. FS: food sensitization. HC: healthy control.

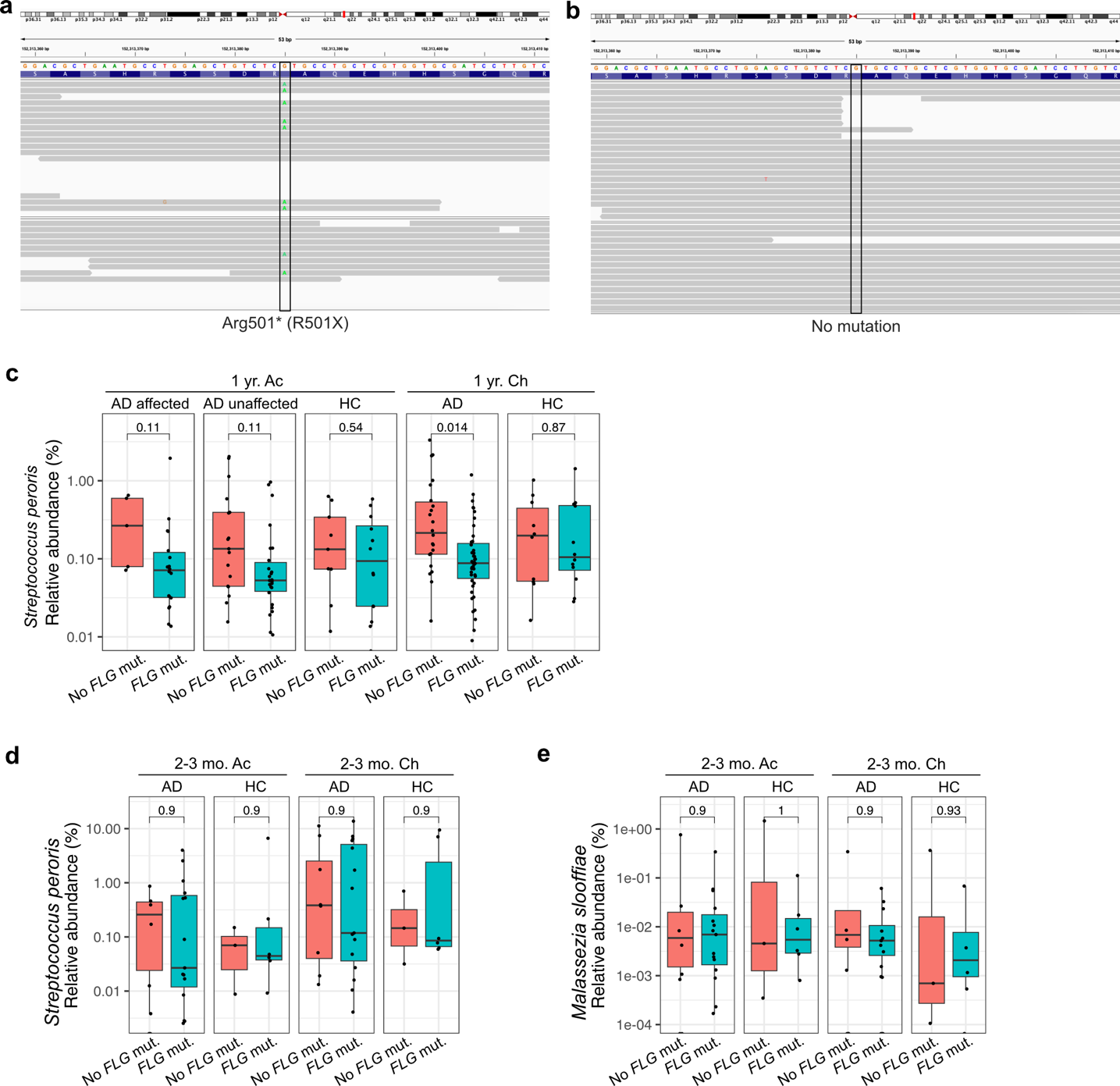

**Figure E4.** (a) Example of an infant with an R501X FLG null mutation and (b) example of an infant without this mutation. (c–d) Relative abundance of Streptococcus peroris (c) and Malassezia slooffiae (d) in the skin of 1-year-old infants, stratified by disease status and by the presence or absence of FLG mutations. (e–g) Relative abundance of Streptococcus mitis (e), Streptococcus peroris (f), and Malassezia slooffiae (g) in the skin of infants at 2–3 months, stratified by disease status and by the presence or absence of FLG mutations. Adjusted p-values from Mann–Whitney U tests are shown. Ac: antecubital fossa. Ch: cheek. AD: atopic dermatitis. FA: food allergy. FS: food sensitization. HC: healthy control. FLG: filaggrin.

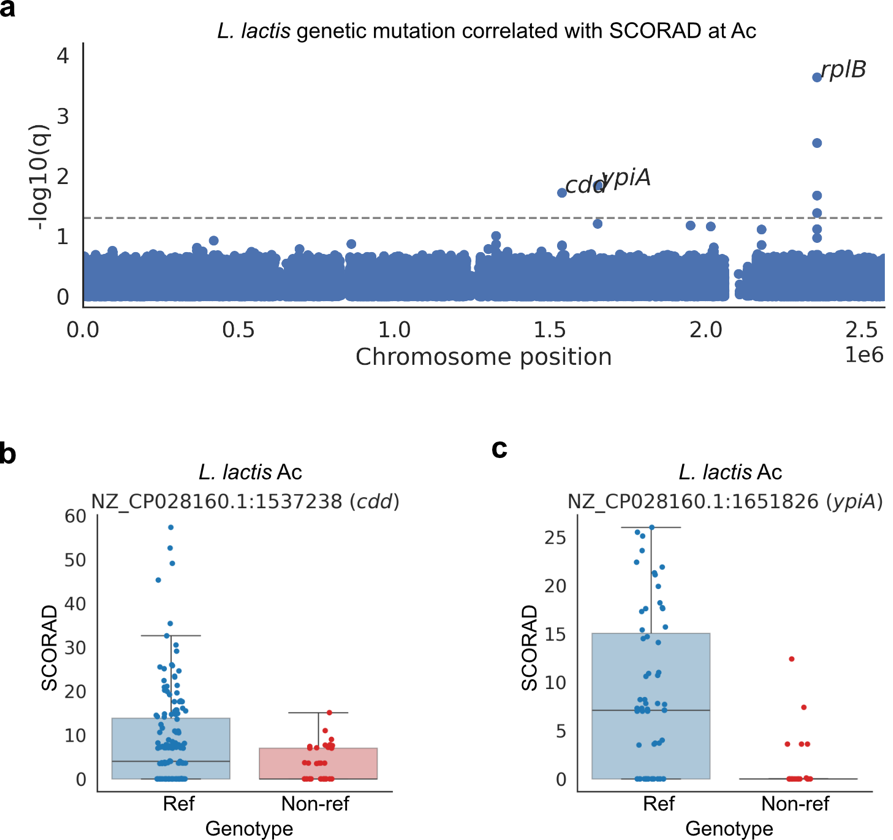

**Figure E5.** (**a**) Manhattan plot showing adjusted p-values for associations between Lactococcus lactis variants and SCORAD scores in antecubital fossa samples. (**b**) Missense mutation in the cdd gene of L. lactis showing a negative association with SCORAD score. (**c**) Missense mutation in the ypiA gene of L. lactis showing a negative association with SCORAD score. Ac: antecubital fossa. Ch: cheek.

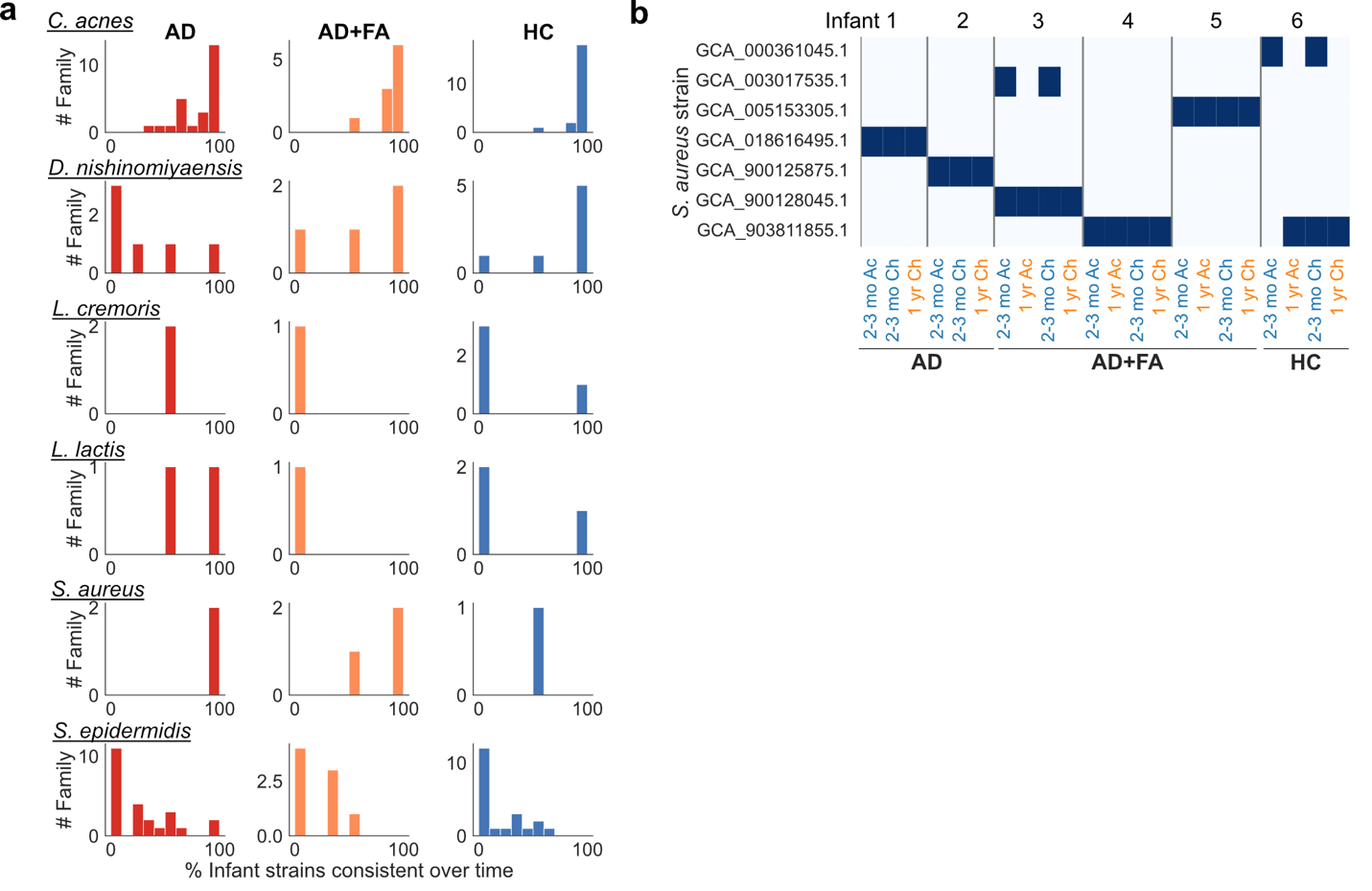

**Figure E6. (a)** Number of families in which strains were detected on infants at both 2–3 months and 1 year, stratified by infant disease status. **(b)** Example of Staphylococcus aureus strain persistence in infants. Grey vertical lines separate families. Ac: antecubital fossa. Ch: cheek. AD: atopic dermatitis. FA: food allergy. HC: healthy control.
